# Complementary evaluation of computational methods for predicting single residue effects on peptide binding specificities

**DOI:** 10.1101/2024.10.18.619108

**Authors:** Merve Ayyildiz, Jakob Noske, Florian J. Gisdon, Josef P. Kynast, Birte Höcker

**Author notes:** equal contribution (Merve Ayyildiz, Jakob Noske, Florian J. Gisdon, Josef P. Kynast).

## Abstract

Understanding the interactions that make up protein-protein or protein-peptide interfaces is a crucial step towards applications in biotechnology. The ability to discriminate between different partners defines the specificity of a binding protein and is equally important as its affinity to the target. Whereas many established computational methods provide an estimate of binding or non-binding, comparing similar ligands is still significantly more challenging. Here we evaluated the capability of predicting ligand binding specificity using three established but conceptually different physics-based methods for protein design. As a model system, we analyzed the binding of peptides to designed armadillo repeat proteins, where a single residue of the peptide was changed systematically, and compared the results with an experimental reference data set. The mutation of a single residue can have a strong impact on binding affinity and specificity, which is difficult to capture in sampling and scoring. We critically assessed the prediction accuracy of the computational methods and found that the prediction performance of each method is differently affected, suggesting the use of a complementary approach of the evaluated methods.

**Author Summary:** Proteins have to recognize other proteins and peptides in the cell with high specificity. To be able to predict such interactions with high precision would be immensely useful for medical and biotechnological applications. Here we tested three computational methods that use physics-based force fields on an experimental dataset and evaluated how well these predictions can be used to discriminate binding pockets on a single residue level. The predicted values of each method and the experimentally determined specificities correlated well, even though each approach had its biases. Therefore, we correlated the predictions with each other to complement the strengths and weaknesses of all approaches.

## Introduction

The function of proteins often relies on the specific recognition of target molecules such as chemical compounds, peptides, or other proteins. This specific recognition is facilitated by a combination of individual interactions at the binding interface. Due to the diverse nature of interactions between amino acids, it is challenging to predict the effect of mutations on the interaction with the target. Affinity towards one target can be reduced or even lost upon mutations at an interface. Specificity, however, does not require to retain the same magnitude of affinity as long as a target is predominantly recognized compared to others. Thus, the design of binding interfaces should not simply focus on the improvement of interactions, but also include the destabilization of undesired complexes [1,2].

Binding affinity and specificity can be determined experimentally; however, this is time- and resource-consuming. Therefore, methods have been developed for the computational evaluation of binding affinities [3,4]. Recently, a number of machine learning approaches have been reported to perform well on predicting protein-protein binding affinity [5–8]. However, due to their dependency on available high-quality experimental data and a less straightforward interpretability, no machine learning model has been as broadly applied as the physics-based methods [9–11].

In this work, we follow a strategy for the evaluation of binding specificities of peptides by combining information obtained from three conceptually different evaluation approaches. First, we used the flex ddG approach [12] from the widely used Rosetta software suite. The flex ddG algorithm calculates binding affinity changes upon mutation. By keeping non-mutated regions fixed, these less critical components of the interaction cancel out which simplifies the calculations. With the backrub approach [13,14], a diverse ensemble is generated, that is structurally close to the initial complex and then is used in the calculation. Second, we applied the Branch and Bound Over K* (BBK*) approach [15] from the Osprey software suite [16]. The BBK* algorithm is based on the approximation of the partition functions for the bound and unbound states of a system, the protein-ligand complex and the free protein and ligand, respectively. The calculated K* scores for a protein-ligand complex approximate the binding affinity constant *K*_a_ and are defined as the quotient of the partition functions for the bound and unbound states [17]. Third, we included our *in-house* tool called PocketOptimizer [18]. PocketOptimizer generates an ensemble of the bound state and determines the energetically best combination of side chain rotamers and the ligand conformation and position in the binding pocket.

As a model system for our study, we chose a designed armadillo repeat protein (dArmRP) with a peptide ligand that is mutated at one distinct position (Fig 1) [19]. The availability of experimental data with a high dynamic range for many single residue variants, where most of the interaction interface stays constant, allows us to focus on a small variable interaction region [20].

**Figure 1.**
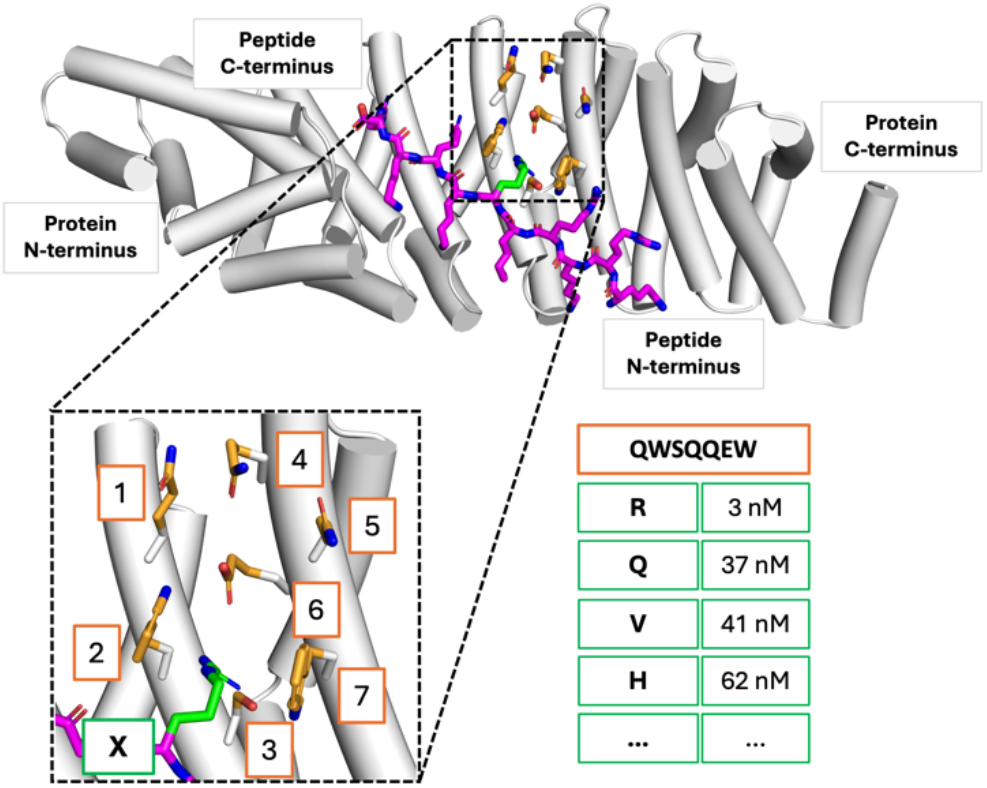
Model system for the computation of single residue effects on peptide binding specificities. The crystal structure of a designed armadillo repeat protein (dArmRP) is shown as white cartoon (PDB ID: 6SA8). A peptide consisting of five alternating Lys and Arg residues ((KR)_5_) that binds to the dArmRP is shown as magenta sticks. The designed binding pocket consisting of seven residues (orange sticks) is shown in the enlarged inset. The side chain of the Arg residue (labelled X) is shown in green. The table illustrates experimental binding affinities for peptides with a mutation of the amino acid that binds to the designed binding pocket (residue identities in the orange box).

We evaluated the relative binding affinities of five experimentally characterized binding pockets to model the binding specificity. For comparison, we correlated these results with the experimentally determined binding affinities. The characteristics and biases of the methods BBK*, flex ddG, and PocketOptimizer are illustrated and demonstrate the benefit of combining complementary results from various computational sources.

## Results and Discussion

We initially applied the three algorithms flex ddG, BBK*, and PocketOptimizer to score protein-peptide complexes where the peptide ligand is mutated at a single position relative to the respective binding pocket of the designed armadillo repeat protein (dArmRP) (Fig 1). By that, we obtained predictions for the binding of all protein-peptide pairs for which experimental binding data was available to evaluate the specificity of the addressed binding pockets. This allowed us to correlate the calculated binding specificities with the experimentally determined ones to asses the quality of the predictions.

### Comparison of binding specificity predictions with experimentally determined data

In a first step we compared the binding specificities calculated using flex ddG, BBK*, and PocketOptimizer with the experimentally determined binding affinities of the so-called Arg-binder (see Fig 1). This was done for calculations based on two different available crystal structures, namely PDB-ID 6SA8 and 5AEI (S1 Fig), thereby gaining insights into the dependency of the predictions on the input structure (Fig 2A and 2B).

**Figure 2.**
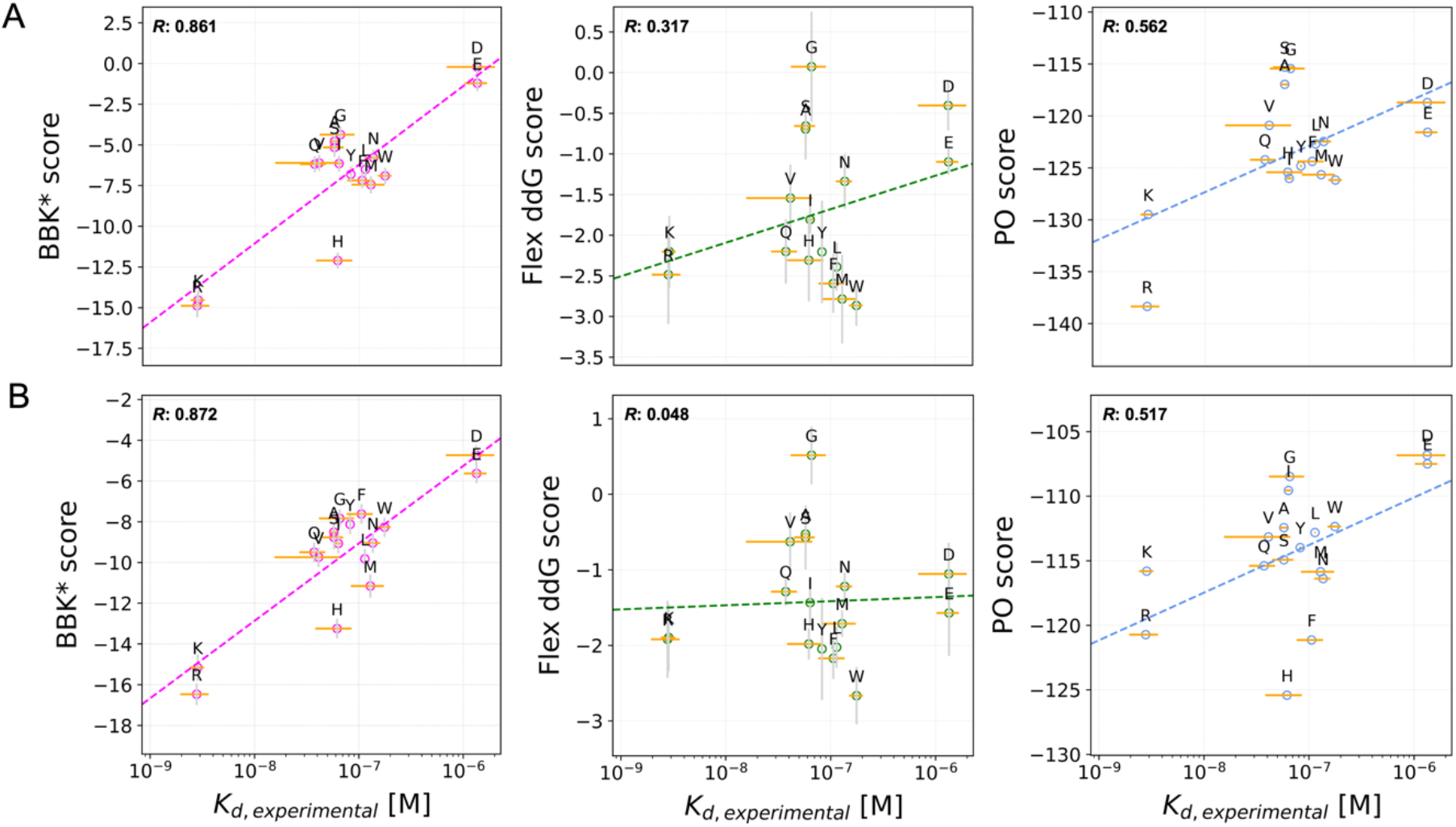
Correlation of calculated and experimentally determined binding specificities. Specificity predictions compared to experimental data for the Arg-binder pocket (Linear fits shown in dashed lines). Binding specificity predictions from BBK* (magenta), flex ddG (green) and PocketOptimizer (blue) were correlated with experimentally determined binding specificities using the crystal structures 6SA8 (A) and 5AEI (B) as scaffold. Pearson correlations are given inside the corresponding plots.

The Arg binding pocket has shown high affinity in experiments for the positively charged amino acids Arg and Lys [21]. The *K*_D_ for these residues is in the low nanomolar range, whereas the *K*_D_ for peptides with the negatively charged amino acids Asp or Glu at the same position is in the micromolar range (Fig 2, *K*_D, *experimental*_). The measurements of peptides with every other proteogenic amino acid at this position are clustered in between with similar *K*_D_ values. The partially overlapping error margins of this middle group add to the similarity and make the individual targets hard to distinguish. Thus, the specificity prediction of BBK* reflected well the experimentally measured trend (Fig 2, left). The calculated specificities allowed to distinguish between the ligands with positively and negatively charged amino acids at the variable position. Ligands with uncharged amino acids showed similar predicted affinities and clustered in between the ligands with positively and negatively charged amino acids at the variable position. Only the amino acid His was predicted to bind with higher affinity compared to the experimental values.

With flex ddG on the other hand, ligands with positively and negatively charged amino acids were distinguished well only when based on the structure 6SA8 (Fig 2A, center), while the difference between these two amino acid groups was not resolved for 5AEI (Fig 2B, center). Small amino acids like Gly, Ala and Ser were predicted to be overall worse than most other amino acids. All other amino acids cluster in the middle with a trend indicative of a bias for large amino acids.

The predictions with PocketOptimizer on the other hand were similar for both scaffolds, though the amino acids Phe and His were emphasized more in the case of 5AEI. Overall, PocketOptimizer distinguished well between the negatively and positively charged amino acids, and even separated Arg from Lys. The calculations based on 6SA8 resulted in overall lower energy scores compared to the scores for 5AEI with negatively charged residues being represented better.

After analyzing a pocket for a charged amino acid, we looked at binding pockets for the aromatic residues Tyr and Trp (Fig 3, first two columns). With a *K*_D_ in the low nanomolar range and about one order of magnitude difference to other measured amino acids, the Tyr binding pocket shows a high affinity and a high specificity for Tyr (Fig 3, *K*_D, *experimental*_). Similarly, the Trp binding pocket shows a tight binding with an experimentally determined *K*_D_ in the nanomolar range and a high specificity towards Trp. Additionally, we analyzed His and Ile binding pockets (Fig 3). Based on the experimental results, the His binding pocket is well able to distinguish His from other amino acids and binds His with a high affinity. The closest amino acid target for this binding pocket is Arg with a *K*_D_ of more than one order of magnitude higher. The binding pocket Ile also shows high affinity for its target; however, the distinction of different amino acids in this position is not as clear since many of the experimentally determined error margins overlap (Fig 3, column 4).

**Figure 3.**
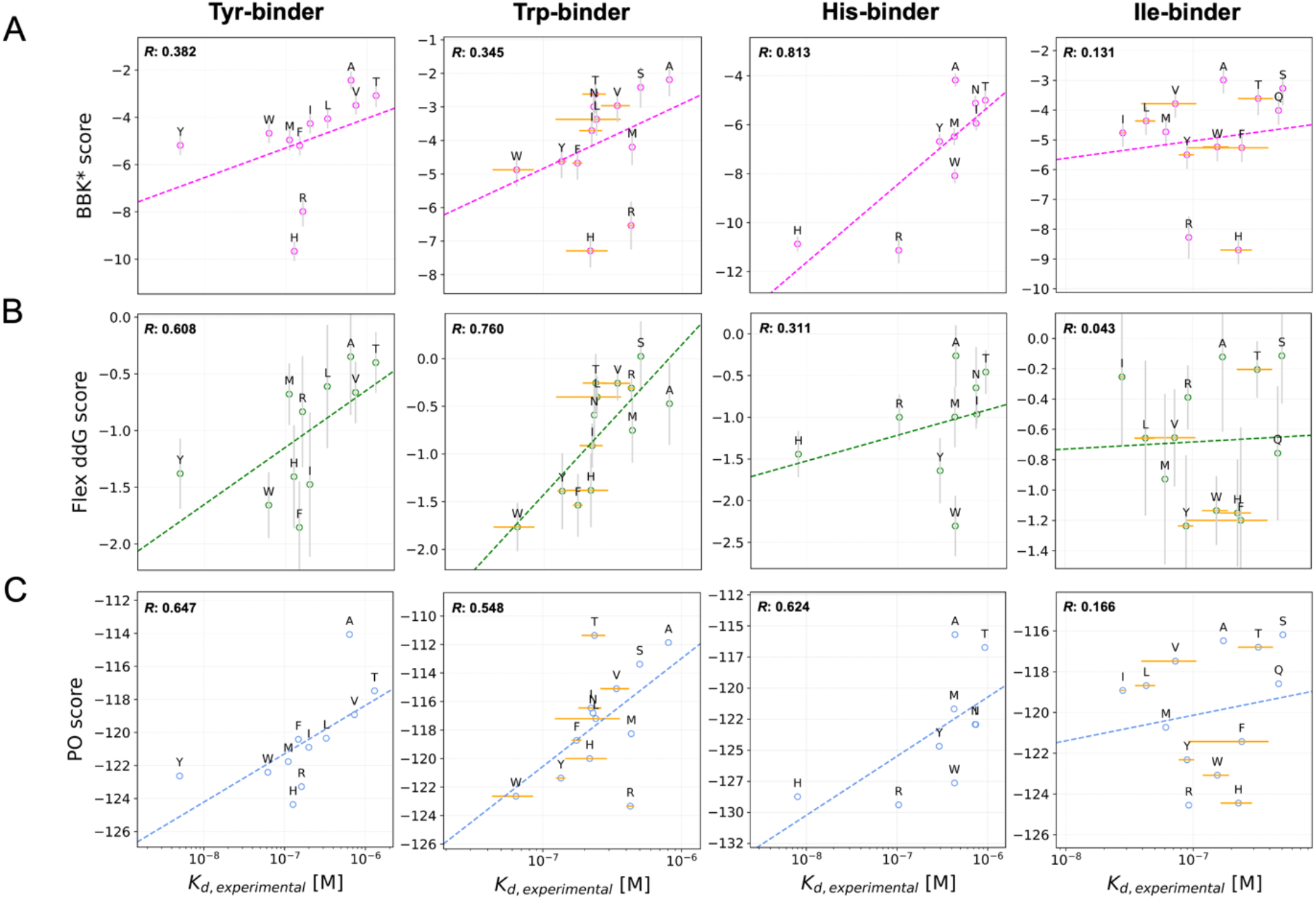
Correlation between specificity predictions and experimental binding data for the Tyr, Trp, His, and Ile binding pockets. Correlation between experimental measurements for each binder with the calculations from BBK* (A), flex ddG (B), and PocketOptimizer (C) using 6SA8 as the scaffold. All predictions are shown with their corresponding Pearson correlations.

The specificity predictions of BBK* for the binding pockets of the aromatic amino acids Tyr and Trp did again recreate the trend observed in the experimental results. Both Tyr and Trp were ranked well for their respective pockets by BBK*, with the exceptions of His and Arg, which were strongly favoured. Both of these binding pockets were also well predicted by flex ddG. For the Trp-binder, all calculated amino acids showed a good correlation with the experimental values. The predictions for the His pocket by BBK* also correlated with the experimental values. In this case His was the second-best prediction for its pocket, with a slight difference from Arg, and both amino acids were predicted significantly better than all other amino acids. In the flex ddG predictions the correlation is not as clear. Although flex ddG can capture the trend of the experimental results for His binding, the amino acids Tyr and Trp were rated with higher affinity indicating a bias for these aromatic amino acids. The specificity calculations by PocketOptimizer on the other hand showed consistently good predictions for Tyr, Trp and His binding pockets, there was however a bias observed towards Arg and His in each case.

In contrast to these generally good correlations, all three methods had difficulties predicting the experimentally observed trends for the Ile binding pocket. While BBK* and PocketOptimizer ranked Ile in the middle of the other amino acids, it is ranked as one of the worst binding ones by the flex ddG method (Fig 3, column 4). For all test cases, Ile was a difficult target to predict as it is not a large or charged amino acid. This also made the Ile binding pocket a more challenging test case. In addition, the experimental values determined for the Ile-binder are closer to each other than for any other binders making the case even more tricky.

Overall, repeating the predictions, which were based on the 5AEI structure, with the 6SA8 structure showed comparable results (S2 Fig). Small differences might be due to the architecture of the constructs. In 6SA8, the peptide-binding interface is protected from crystal contacts by a fused DARPin protein, while the binding interface of the crystal structure 5AEI is not protected and is involved in some crystal contacts (S1 Fig). However, the DARPin fusion might also affect the bending shape of the armadillo repeat super helix, which could also lead to slight changes in the prediction results.

Since the predictions with the different scaffold structures were comparable, we focused on predicting binding specificities by comparing different targets for one binder with a defined binding pocket modelled from one scaffold. It should be noted though that the absolute values were not comparable between the different binding pockets or scaffolds.

### Prediction method biases

There are known tendencies towards overpredicting energetic contributions of certain amino acids [22,23]. We observed similar biases in our predictions. To address individual tendencies of the methods towards different amino acid types, we calculated a relative bias for each target peptide with the corresponding amino acid (Fig 4). A trend was observed towards an overemphasis of larger amino acids which hints at the tendency towards a higher energetic value for larger amino acids. Prediction offsets for Tyr were lower on average, but the individual offsets were spread throughout the offset range. This might be due to Tyr having a high affinity to most of the examined binding pockets, also the calculation of the offsets was less accurate for the edge cases. For charged amino acids, which are medium to large, the prediction offset was low, especially for Lys, Glu and Asp. However, these three amino acids lack experimental validation for several pockets making the data basis rather small. Further, Arg is highly overemphasized by BBK* and PocketOptimizer. In comparison the flex ddG predictions seemed more balanced. Despite these trends the calculation of relative offsets needs to be interpreted with caution; due to the normalization of the offset, its absolute error might differ drastically for different pockets.

**Figure 4.**
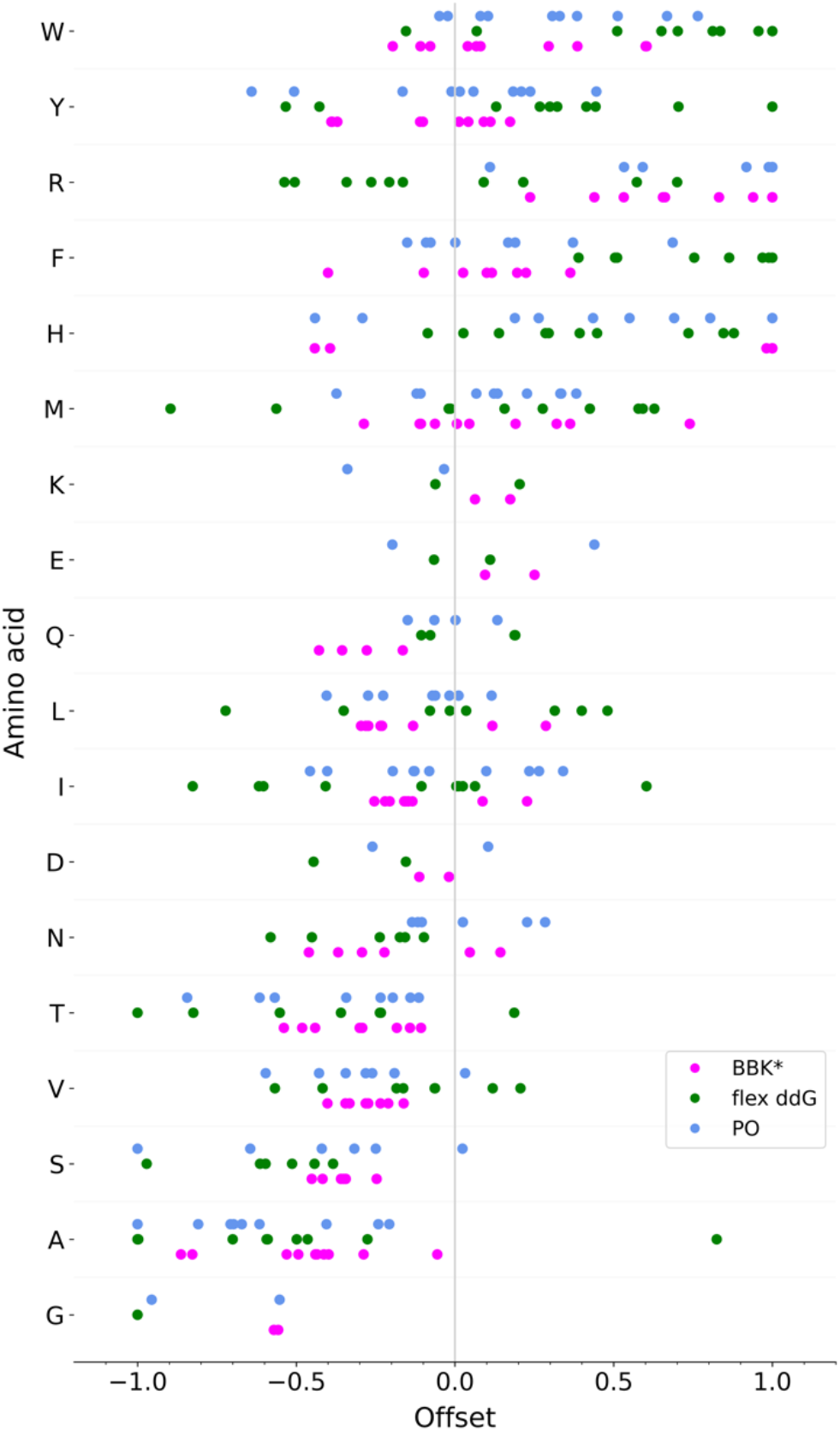
Individual relative offsets from optimal fit for individual amino acid targets. Amino acids are listed at the y-axis according to their relative mass.

**Figure 5:**
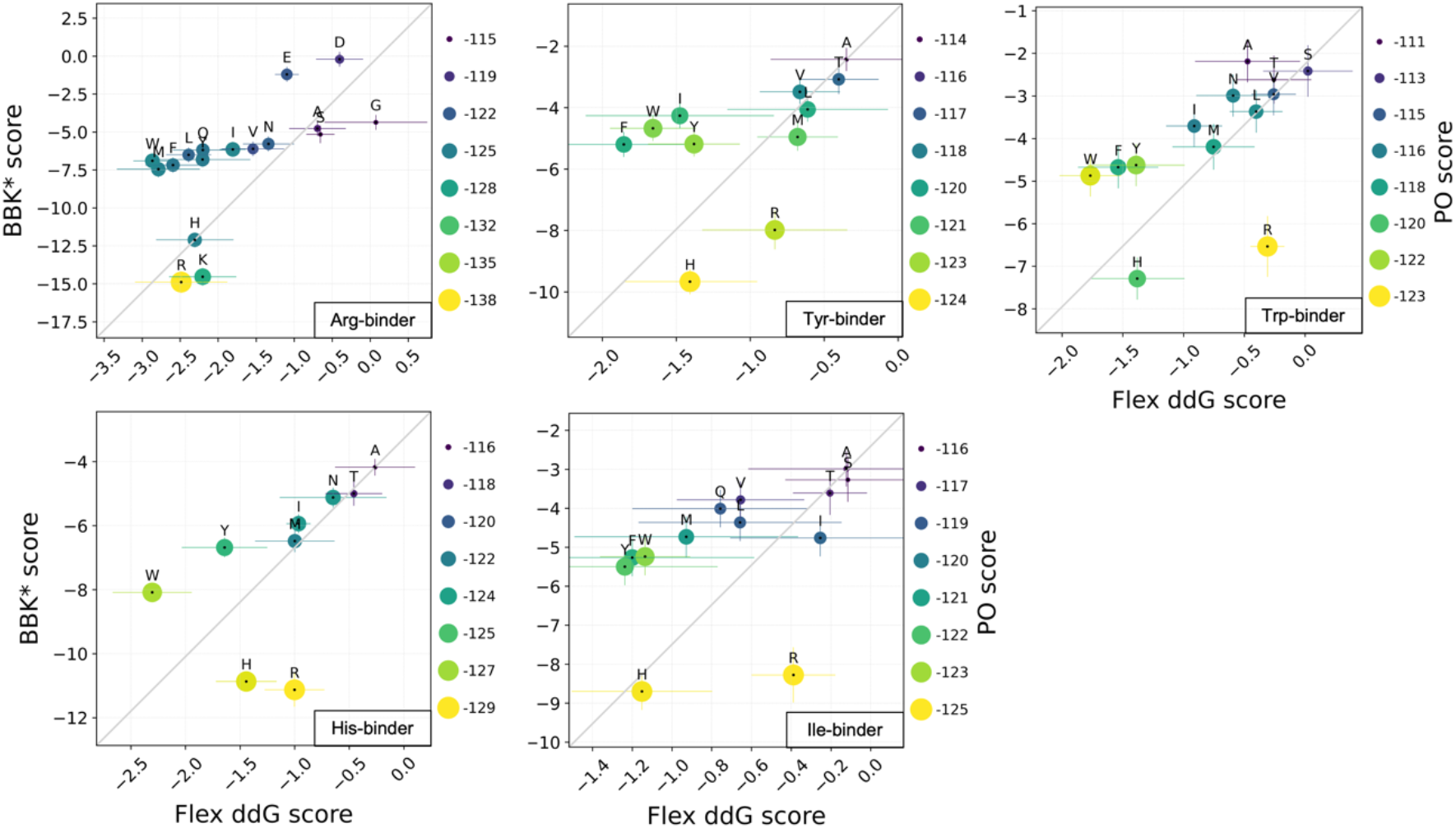
Correlation of specificity predictions from all three methods. BBK*, flex ddG, and PocketOptimizer predictions for Arg, Tyr, Trp, His and Ile binders were obtained using the crystal structure 6SA8 as the scaffold. Flex ddg and BBK* values are shown on the x and y axes, respectively, while PocketOptimizer values are shown as color and size of the data points with the values written on the right side of each plot. Error bars are from flex ddG and BBK* calculations.

### Correlations between binding specificity predictions from BBK*, flex ddG and PocketOptimizer

As we could observe general tendencies for the predictions of some amino acids in our calculations, we correlated the methods BBK*, flex ddG, and PocketOptimizer with each other considering the five binders analyzed above. The aim was to analyze the strengths that each method has and increase the accuracy of specificity predictions by combining results from different methods. For the Arg-binder, BBK* and PocketOptimizer values correlated well with each other. For the Tyr- and Trp-binders flex ddG performed better and corrected His and Arg tendencies that the other two methods show. Since the Ile-binder was not predicted well by any of the methods, while the His-binder was predicted well by all, the correlation was less informative for these two binders. Nonetheless, the correlations are helpful by providing robustness of the predictions. A similar overall trend could also be seen for predictions based on scaffold 5AEI (S3 Fig)

## Conclusion

Peptide-binding pocket designs were evaluated with complementary binding specificity calculations using three conceptually different methods. BBK* from Osprey as well as PocketOptimizer rely on fully physics-based force fields whereas flex ddG from Rosetta also includes empirical terms for its energy function. We compared the specificities of single amino acid binding pockets for different residue types and evaluated the prediction accuracy of the three approaches. Flex ddG seemed to more accurately predict the specificity of binding pockets for aromatic amino acids, while BBK* performed well on pockets for charged amino acids. PocketOptimizer on the other hand showed a lower, but more consistent Pearson correlation in those test cases. These shortcomings can be accounted for by a complementary analysis using all three methods in combination. This way, individual biases can be corrected by another prediction method. Besides, knowing about the different tendencies of each method, prediction accuracies can be improved.

With this evaluation, we provide a detailed overview of the prediction accuracy of widely used physics-based protein design software for binding specificity. It is apparent that single residue predictions are still a great challenge and need to be addressed in the field. There is hope that this can be tackled by new approaches, such as machine learning-based algorithms. The analysis and benchmark provided here will be a valuable instrument to test and evaluate their prediction accuracy.

## Materials and Methods

### Preparation of structural models

dArmRPs generated through consensus design [24] were chosen to serve as scaffold proteins since they selectively bind extended peptides in a modular fashion. The proteins bind to a repetitive, positively charged sequence, namely a (KR)_5_ peptide. All simulations were based on the two crystallographic structures with PDB-ID 6SA8 and 5AEI that were both solved in complex with the (KR)_5_ ligand. The crystal structure 6SA8 entails a dArmRP-fusion with a designed ankyrin repeat protein (DARPin) that arranges in a ring-like structure [25] while 5AEI includes only the dArmRP [21]. The structures were prepared for the simulations with MoleculeKit [26] as follows: All ions, water molecules and crystallization additives were removed. For the models based on 6SA8, the protein residues 179 to 514 of chain A were used so that the DARPin fusion was excluded. Chain B with residues 1 to 10 was used as ligand. For the models based on 5AEI, the protein residues 11 to 291 of chain A were taken. The residues 1 to 10 of chain D were used as ligand.

For each structure, 5AEI and 6SA8, five models were prepared according to experimentally characterized protein sequences for an Arg-binder, a Tyr-binder, a Trp-binder, a His-binder, and an Ile-binder [20]. All mutations were introduced with PyMOL [27] and 2000 cycles of molecular sculpting were performed, which is a procedure within PyMOL to return local atomic geometries to the initial configuration of the crystal structure. Seven residues in the binding pockets of the model proteins were mutated according to experimentally determined sequences [20]. The respective residue numbers for models based on 6SA8 are 364, 368, 371, 403, 406, 407, and 410, while for models based on 5AEI, they are 155, 159, 162, 194, 197, 198, and 201.

In the 6SA8 structure, residue 6 of the peptide ligand binds to the prepared binding pocket. For the calculations with BBK* and flex ddG, residue 6 for the input structure was modeled as alanine and was allowed to mutate to all other amino acids within the respective method. For the calculations with PocketOptimizer, all input structures with the respective amino acid at position 6 of the ligand were generated. In accordance with the experimental setup, residue 4 and 8 of the ligand were modeled as alanines. In 5AEI, the respective residue of the peptide ligand is residue 4 (S1 Fig).

Protonation states were determined with PropKa 3.2 [28]. Uncertain protonation states of amino acids, which were predicted to titrate less than 0.5 pH units from the adjusted pH 7, were manually adjusted according to present interactions.

### Computational evaluation of binding affinity scores

For the evaluation of binding affinities, we used three different methods: (1) BBK* in Osprey 3.2.304 [15], (2) flex ddG in Rosetta 3.12 [12], and (3) PocketOptimizer 2.0 [18].

1. For the calculations with the BBK* algorithm, we applied side chain flexibility to all seven residues, which specify the binding pocket (Fig 1). Four of the seven residues were modeled with continuous flexibility, which is the option to rotate continuously within a range of ±9° from the discrete values of the rotamer library [29]. For models that are based on 6SA8, discrete flexibility was applied to residues 364, 403, and 406, and continuous flexibility to residues 368, 371, 407, and 410. For the ligand, residue 6 was modeled with continuous flexibility. Backbone flexibility was applied to ligand residues 2 to 8. For the models that are based on 5AEI, discrete flexibility was applied to residues 155, 194, and 197 and continuous flexibility to the residues 159, 162, 198 and 201. For the ligand, residue 4 was modeled with continuous flexibility. Backbone flexibility was applied to ligand residues 2 to 8. For the calculations, we used the implemented Amber96 forcefield with the EEF1 solvation model [30]. We applied an epsilon of 0.68 as suggested to be sufficient to distinguish between orders of magnitudes of K*.
2. For the simulations with the flex ddG algorithm, we used the same protocol as described in [12]. For each model, we generated ensembles with 250 structures. A sufficient ensemble size for robust results was determined with initial tests. For each generated structure, the backrub algorithm [14] was run for 35000 backrub Monte Carlo steps. A snapshot structure was stored every 5000 steps, resulting in seven structures. We used the Talaris all-atom energy function [31]. The score analysis was performed as described, using the corresponding reweighting scheme based on a generalized additive model [12].
3. The program PocketOptimizer 2.0 [18] was used with the Amber ff14SB force field [32] to calculate all interaction energies between flexible side chains, ligand poses and the fixed scaffold structure. Since the dArmRP is highly charged, the electrostatic component was scaled down to 1 % based on initial tests. Minimization of side chains was carried out by applying a constant force of 5 kcal·mol^-1^ to all heavy backbone atoms and the energy was minimized until convergence. Rotamer sampling for all pocket positions was carried out, selecting rotamers from the Dunbrack backbone-dependent rotamer library with a probability of 1 % or above [33]. For the mutatable ligand position the C.M. Lib backbone-independent rotamer library [34] was used. All sampled peptide conformations were translated by ± 0.1 Å and rotated by ± 2.5° along each axis. Of the rotamers and peptide poses sampled, those with a vdW energy of more than 50 kcal·mol^-1^ in a scaffold where all pocket positions are mutated to an alanine were pruned before the energy calculations.

### Processing and analysis of calculated binding affinity scores

The output data of BBK* and flex ddG was further processed. For our analysis, we restricted the comparison of the predicted scores to the available experimental data (S1 Table). The obtained K* scores from BBK* were converted into approximate p*K*_D_ values. As an uncertainty range, the obtained upper and lower bounds of the K* score were taken. The obtained flex ddG scores represented binding affinity changes in kcal·mol^-1^ relative to the respective alanine references. We considered the best 5% of the calculated values from the simulated ensemble and calculated the mean value with the mean absolute error to represent the average score for the energetic change in the binding energy. In PocketOptimizer the scores represent the binding energy in kcal·mol^-1^ between the protein and the peptide. Since PocketOptimizer selects one combination of pocket rotamers and a single ligand pose that represent the global minimum energy conformation (GMEC) in the sampled set of conformations, there is only a single energy value without error computed for each complex.

### Calculation of prediction biases

For each method on each tested binding complex, a linear fit was calculated between the experimentally determined binding affinity and the predicted scores. The offset of the predicted score from each fit was extracted and normalized to limit the highest absolute offset for each method/pocket combination to 1. This way, we extracted individual relative offsets from an optimal fit for individual amino acid targets. We sorted the relative offsets in the order of the molecular mass of the amino acid.

## Acknowledgments

We thank Emily Freund for testing our prediction protocols and acknowledge Bruce Donald and his lab members for support on using OSPREY. This work was supported by European Union Horizon 2020 FETOPEN program PRe-ART, grant agreement No. 764434, and European Union EIC Transition program PRe-ART-2T, grant agreement No. 101058027.

## Supplementary Information

### Supplementary figures 1-3

**Figure S1:**
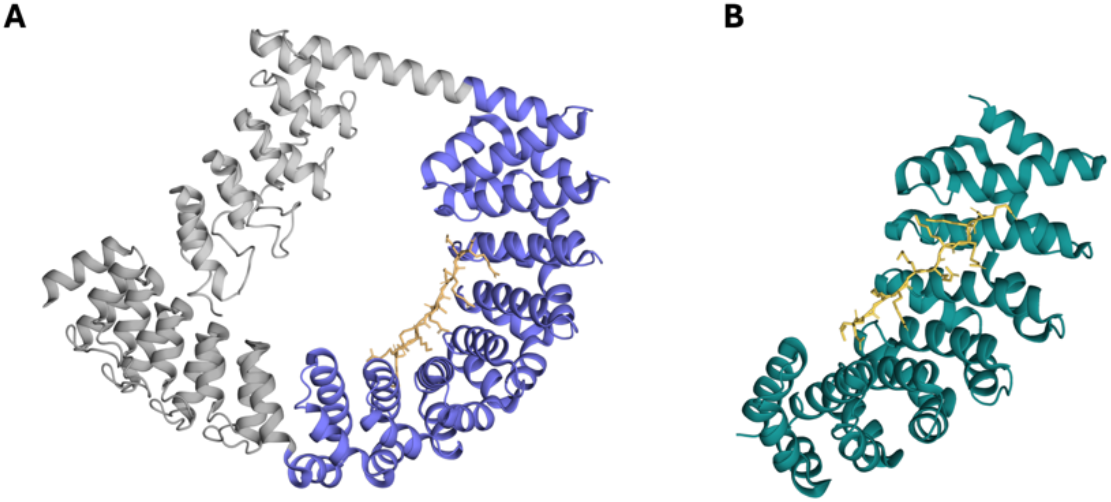
Crystal structures of dArmRP proteins used in this study. The structures with PDB-ID 6SA8 (A) and 5AEI (B), both with bound (KR)_5_ peptides shown as orange sticks, were used for the calculations.

**Figure S2:**
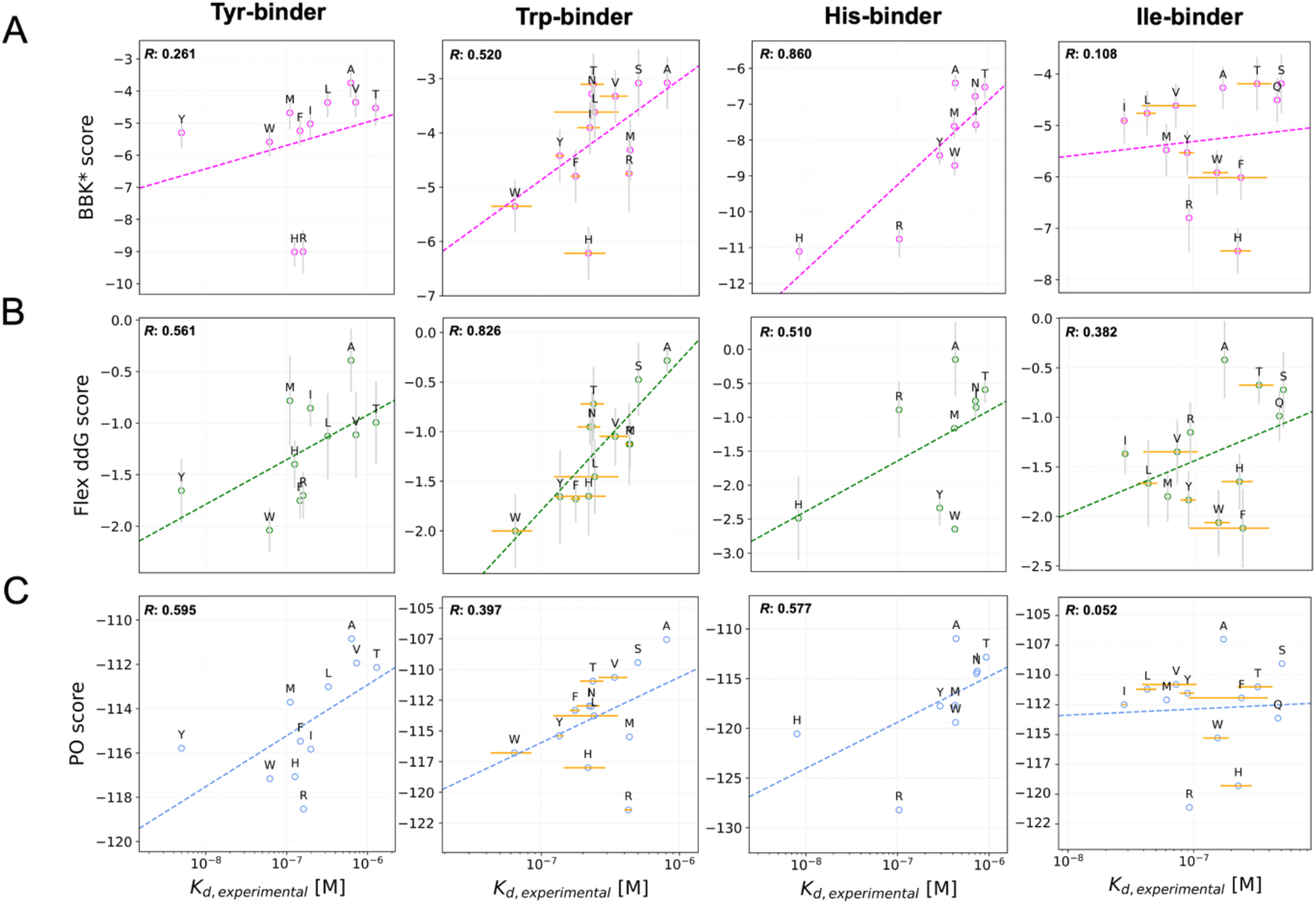
Correlation between predicted and experimentally determined binding specificities. Crystal structure 5AEI was used as scaffold where binding pockets for Tyr, Trp, His, and Ile were introduced.

**Figure S3:**
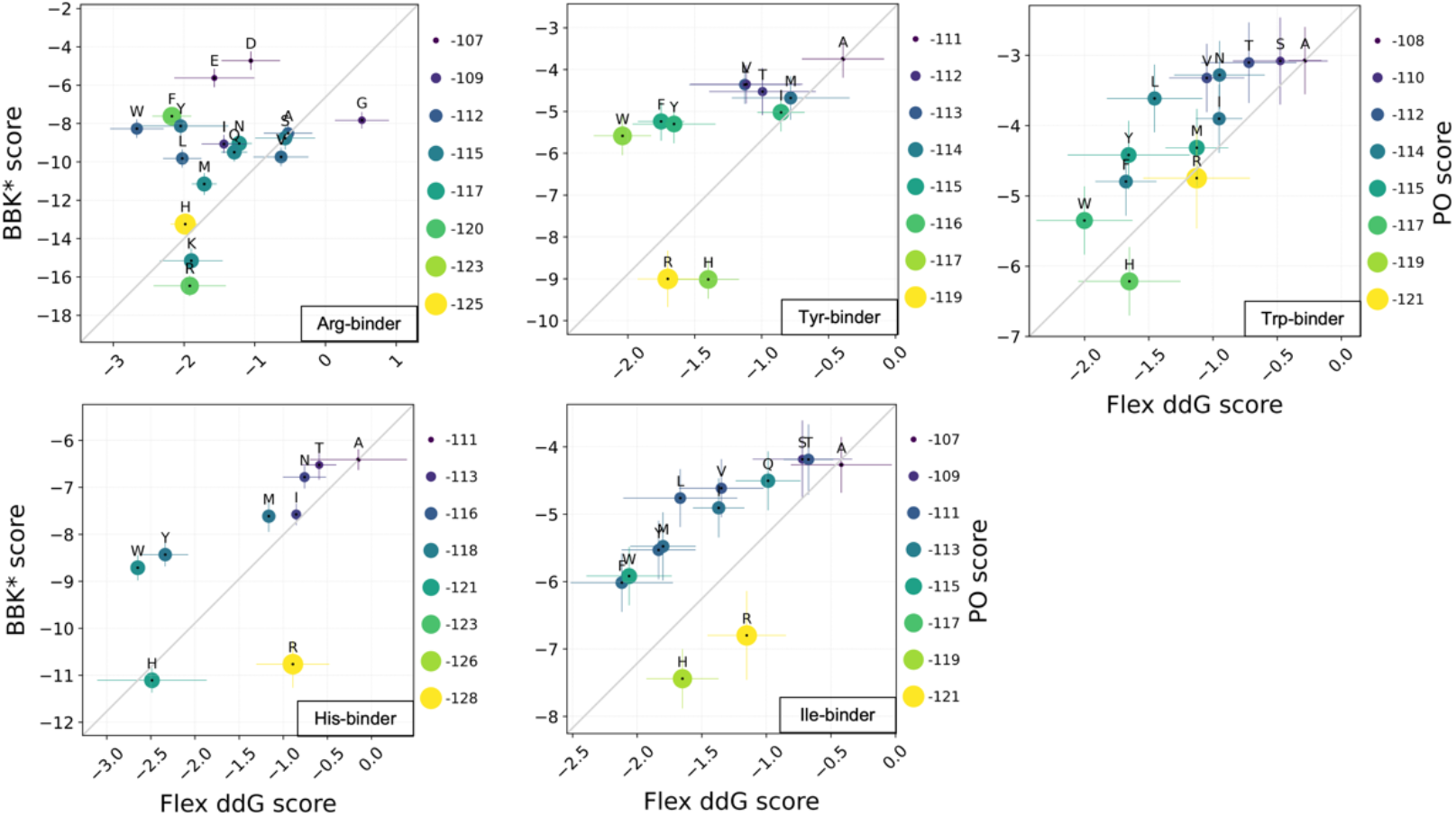
Correlation of calculated binding specificity predictions. Crystal structure 5AEI was used as scaffold for the introduction of Arg, Tyr, Trp, His and Ile binding pockets and the algorithms flex ddG, BBK* and PocketOptimizer were used for the specificity predictions.

### Supplementary table 1

**Table S1:** Experimental binding affinity data used for comparison with calculated scores.

| **Arg-binder (QWSQQEW)** |  |  |
| --- | --- | --- |
| Peptide | $K_D$ [nM] | Error [nM] |
| R | 2.787 | 0.85 |
| K | 2.85 | 0.44 |
| Q | 37.245 | 10.37 |
| V | 41.055 | 25.52 |
| A | 57.57 | 4.79 |
| S | 57.64 | 13.0 |
| H | 61.8 | 23.48 |
| I | 63.63 | 4.14 |
| G | 65.665 | 24.19 |
| Y | 82.52 | 1.94 |
| F | 105.92 | 29.81 |
| L | 113.95 | 1.63 |
| M | 128.64 | 44.92 |
| N | 136.1 | 23.90 |
| W | 176.3 | 26.45 |
| D | 1334 | 651.95 |
| E | 1343.5 | 323.15 |

| **Tyr-binder (KEVLIRQ)** |  |  |
| --- | --- | --- |
| Peptide | $K_D$ [nM] | Error [nM] |
| Y | 5 | - |
| W | 62 | - |
| M | 111 | - |
| H | 127 | - |
| F | 148 | - |
| R | 162 | - |
| I | 199 | - |
| L | 327 | - |
| A | 636 | - |
| V | 731 | - |
| T | 1296 | - |

| **Trp-binder (TATAWRT)** |  |  |
| --- | --- | --- |
| Peptide | $K_D$ [nM] | Error [nM] |
| W | 64 | 21 |
| Y | 135 | 10.68 |
| F | 175.5 | 14.5 |
| H | 217.5 | 72.5 |
| I | 222.5 | 42.12 |
| N | 230 | - |
| T | 236 | 46 |
| L | 241 | 119 |
| V | 339.5 | 79.5 |
| R | 427.5 | 27.5 |
| M | 434.5 | - |
| S | 500 | - |
| A | 806 | - |

| **Ile-binder (FALYDRV)** |  |  |
| --- | --- | --- |
| Peptide | $K_D$ [nM] | Error [nM] |
| I | 28 | 1 |
| L | 42.75 | 7.75 |
| M | 61 | - |
| V | 72.5 | 33.5 |
| R | 92.5 | - |
| Y | 89.5 | 12.4 |
| W | 155 | 36 |
| A | 172.9 | - |
| H | 227 | 63 |
| F | 241 | 150 |
| T | 324.7 | 100.21 |
| Q | 470 | - |
| S | 507 | - |

| His-binder (DYTDWQA) |  |  |
| --- | --- | --- |
| Peptide | $K_D$ [nM] | Error [nM] |
| H | 8 | - |
| R | 104 | - |
| Y | 292 | - |
| M | 425 | - |
| W | 430 | - |
| A | 436 | - |
| N | 727 | - |
| I | 740 | - |
| T | 930 | - |

## References

1. Beard H, Cholleti A, Pearlman D, Sherman W, Loving KA. Applying physics-based scoring to calculate free energies of binding for single amino acid mutations in protein-protein complexes. PLoS One. 2013;8: 1–11. doi:10.1371/journal.pone.0082849

2. Sharabi O, Shirian J, Shifman JM. Predicting affinity- and specificity-enhancing mutations at protein-protein interfaces. Biochem Soc Trans. 2013;41: 1166–1169. doi:10.1042/BST20130121

3. Siebenmorgen T, Zacharias M. Computational prediction of protein–protein binding affinities. Wiley Interdiscip Rev Comput Mol Sci. 2020;10: 1–18. doi:10.1002/wcms.1448

4. Dutkiewicz Z. Computational methods for calculation of protein-ligand binding affinities in structure-based drug design. Phys Sci Rev. 2022;7: 933–968. doi:10.1515/psr-2020-0034

5. Liu X, Luo Y, Li P, Song S, Peng J. Deep geometric representations for modeling effects of mutations on protein-protein binding affinity. PLoS Comput Biol. 2021;17: 1–28. doi:10.1371/journal.pcbi.1009284

6. Liu X, Feng H, Wu J, Xia K. Persistent spectral hypergraph based machine learning (PSH-ML) for protein-ligand binding affinity prediction. Brief Bioinform. 2021;22: 1–12. doi:10.1093/bib/bbab127

7. Casadio R, Martelli PL, Savojardo C. Machine learning solutions for predicting protein– protein interactions. Wiley Interdiscip Rev Comput Mol Sci. 2022;12. doi:10.1002/wcms.1618

8. Mohseni Behbahani Y, Laine E, Carbone A. Deep Local Analysis deconstructs protein-protein interfaces and accurately estimates binding affinity changes upon mutation. Bioinformatics. 2023;39: I544–I552. doi:10.1093/bioinformatics/btad231

9. Guo Z, Yamaguchi R. Machine learning methods for protein-protein binding affinity prediction in protein design. Front Bioinforma. 2022;2: 1–11. doi:10.3389/fbinf.2022.1065703

10. Tang T, Zhang X, Liu Y, Peng H, Zheng B, Yin Y, et al. Machine learning on protein–protein interaction prediction: models, challenges and trends. Brief Bioinform. 2023;24: 1–11. doi:10.1093/bib/bbad076

11. Lannelongue L, Inouye M. Pitfalls of machine learning models for protein–protein interaction networks. Bioinformatics. 2024;40. doi:10.1093/bioinformatics/btae012

12. Barlow KA, Ó Conchúir S, Thompson S, Suresh P, Lucas JE, Heinonen M, et al. Flex ddG: Rosetta Ensemble-Based Estimation of Changes in Protein-Protein Binding Affinity upon Mutation. J Phys Chem B. 2018;122: 5389–5399. doi:10.1021/acs.jpcb.7b11367

13. Davis IW, Arendall WB, Richardson DC, Richardson JS. The backrub motion: How protein backbone shrugs when a sidechain dances. Structure. 2006;14: 265–274. doi:10.1016/j.str.2005.10.007

14. Friedland GD, Linares AJ, Smith CA, Kortemme T. A simple model of backbone flexibility improves modeling of side-chain conformational variability. J Mol Biol. 2008;380: 757– 774. doi:10.1016/j.jmb.2008.05.006

15. Ojewole AA, Jou JD, Fowler VG, Donald BR. BBK* (Branch and Bound over K*): A Provable and Efficient Ensemble-Based Protein Design Algorithm to Optimize Stability and Binding Affinity over Large Sequence Spaces. J Comput Biol. 2018;25: 726–739. doi:10.1089/cmb.2017.0267

16. Hallen MA, Martin JW, Ojewole A, Jou JD, Lowegard AU, Frenkel MS, et al. OSPREY 3.0: Open-source protein redesign for you, with powerful new features. J Comput Chem. 2018;39: 2494–2507. doi:10.1002/jcc.25522

17. Lilien RH, Stevens BW, Anderson AC, Donald BR. Algorithm for Protein Redesign and Its Application Synthetase A Phenylalanine Adenylation Enzyme. J Comput Biol. 2005;12: 740–761.

18. Noske J, Kynast JP, Lemm D, Schmidt S, Höcker B. PocketOptimizer 2.0: A modular framework for computer-aided ligand-binding design. Protein Sci. 2023;32: 1–8. doi:10.1002/pro.4516

19. Gisdon FJ, Kynast JP, Ayyildiz M, Hine A V., Plückthun A, Höcker B. Modular peptide binders-development of a predictive technology as alternative for reagent antibodies. Biol Chem. 2022;403: 535–543. doi:10.1515/hsz-2021-0384

20. Stark Y, Menard F, Jeliazkov JR, Ernst P, Chembath A, Ashraf M, et al. Modular binder technology by NGS-aided, high-resolution selection in yeast of designed armadillo modules. Proc Natl Acad Sci. 2024;121: e2318198121. doi:10.1073/pnas.2318198121

21. Hansen S, Tremmel D, Madhurantakam C, Reichen C, Mittl PRE, Plückthun A. Structure and Energetic Contributions of a Designed Modular Peptide-Binding Protein with Picomolar Affinity. J Am Chem Soc. 2016;138: 3526–3532. doi:10.1021/jacs.6b00099

22. Usmanova DR, Bogatyreva NS, Bernad JA, Eremina AA, Gorshkova AA, Kanevskiy GM, et al. Self-consistency test reveals systematic bias in programs for prediction change of stability upon mutation. Bioinformatics. 2018;34: 3653–3658. doi:10.1093/bioinformatics/bty340

23. Tsishyn M, Pucci F, Rooman M. Quantification of biases in predictions of protein – protein. 2024;25: 1–11.

24. Parmeggiani F, Pellarin R, Larsen AP, Varadamsetty G, Stumpp MT, Zerbe O, et al. Designed Armadillo Repeat Proteins as General Peptide-Binding Scaffolds: Consensus Design and Computational Optimization of the Hydrophobic Core. J Mol Biol. 2008;376: 1282–1304. doi:10.1016/j.jmb.2007.12.014

25. Ernst P, Honegger A, van der Valk F, Ewald C, Mittl PRE, Plückthun A. Rigid fusions of designed helical repeat binding proteins efficiently protect a binding surface from crystal contacts. Sci Rep. 2019;9: 1–10. doi:10.1038/s41598-019-52121-9

26. Doerr S, Harvey MJ, Noé F, De Fabritiis G. HTMD: High-Throughput Molecular Dynamics for Molecular Discovery. J Chem Theory Comput. 2016;12: 1845–1852. doi:10.1021/acs.jctc.6b00049

27. Schrödinger LLC. The PyMol Molecular Graphics System. 2015;2.5.0.

28. Olsson MHM, SØndergaard CR, Rostkowski M, Jensen JH. PROPKA3: Consistent treatment of internal and surface residues in empirical p K a predictions. J Chem Theory Comput. 2011;7: 525–537. doi:10.1021/ct100578z

29. Gainza P, Roberts KE, Donald BR. Protein design using continuous rotamers. PLoS Comput Biol. 2012;8. doi:10.1371/journal.pcbi.1002335

30. Lazaridis T. Effective energy function for proteins in lipid membranes. Proteins Struct Funct Genet. 2003;52: 176–192. doi:10.1002/prot.10410

31. O’Meara MJ, Leaver-Fay A, Tyka MD, Stein A, Houlihan K, Dimaio F, et al. Combined covalent-electrostatic model of hydrogen bonding improves structure prediction with Rosetta. J Chem Theory Comput. 2015;11: 609–622. doi:10.1021/ct500864r

32. Maier JA, Martinez C, Kasavajhala K, Wickstrom L, Hauser KE, Simmerling C. ff14SB: Improving the Accuracy of Protein Side Chain and Backbone Parameters from ff99SB. J Chem Theory Comput. 2015;11: 3696–3713. doi:10.1021/acs.jctc.5b00255

33. Shapovalov M V., Dunbrack RL. A smoothed backbone-dependent rotamer library for proteins derived from adaptive kernel density estimates and regressions. Structure. 2011;19: 844–858. doi:10.1016/j.str.2011.03.019

34. Malisi C, Schumann M, Toussaint NC, Kageyama J, Kohlbacher O, Höcker B. Binding Pocket Optimization by Computational Protein Design. PLoS One. 2012;7. doi:10.1371/journal.pone.0052505

